# Discovery of a prevalent picornavirus by visualizing zebrafish immune responses

**DOI:** 10.1101/823989

**Authors:** Keir M. Balla, Marlen C. Rice, James A. Gagnon, Nels C. Elde

## Abstract

The discovery of new viruses currently outpaces our capacity for experimental examination of infection biology. To better couple virus discovery with immunology, we genetically modified zebrafish to visually report on virus infections. After generating a strain that expresses green fluorescent protein (GFP) under an interferon-stimulated gene promoter, we repeatedly observed transgenic larvae spontaneously expressing GFP days after hatching. RNA sequencing of GFP-positive zebrafish revealed a picornavirus distantly related to known viruses. We detected this virus in publicly available sequencing data from seemingly asymptomatic zebrafish in research institutes world-wide. These data revealed widespread tissue tropism and identified the clonal CG2 strain as notably vulnerable to picornavirus infection. Infection induces the expression of numerous antiviral immune defense genes in zebrafish, an effect also evident in public data. Our study describes a prevalent picornavirus that infects zebrafish, and provides new strategies for simultaneously discovering viruses and their impact on vertebrate hosts.

## Introduction

Viruses are estimated to outnumber their cellular hosts in most environments (Cobian Guemes et al., 2016; Suttle, 2005), indicating that animals regularly encounter and interact with viruses. The ubiquity of viruses is demonstrated by the widespread success of virus discoveries made through metagenomic sequencing of animal feces and tissues (Li et al., 2010; Shi et al., 2018, 2016). While surveys of virus prevalence in animals can reveal close associations, most metagenomic virus discovery studies are agnostic as to how, or even if, the virus and animal interact. Metagenomic investigations of animals with undiagnosed disease have resulted in discoveries of new viruses as the causal agents (Finkbeiner et al., 2008; Greninger et al., 2009; Lysholm et al., 2012), thereby establishing a particularly strong link between virus and host. However, the high frequency at which viruses are discovered in asymptomatic hosts suggests that extreme disease phenotypes result from only a small fraction of virus-host interactions.

Most consequential interactions between viruses and animals involve host immune responses. Thus, one sensitive strategy for discovering previously unknown viruses with clear host interactions is to select for engagement of antiviral immune responses instead of overt disease. While typically invisible, antiviral immune responses can be visualized in some live hosts with the genetic tools that are available for a growing collection of experimentally accessible organisms (Lienenklaus et al., 2009; Palha et al., 2013). Furthermore, carrying out virus discovery in research organisms holds the potential for developing new tractable systems for studying host-virus interactions, as demonstrated through metagenomics approaches in worms, flies, and mice (Felix et al., 2011; Roediger et al., 2018; Webster et al., 2015).

Virus discovery in vertebrates has predominantly been carried out in mammals (Olival et al., 2017), but recent wide-ranging metagenomic surveys revealed that fish are infected by an extensively overlapping set of diverse virus families (Geoghegan et al., 2018; Shi et al., 2018). Viruses discovered in fish can therefore illuminate facets of virus and host biology that are shared across a much wider interval of evolutionary history than traditionally considered. Additionally, increasing our knowledge of the viruses that infect fish is an important goal for improving marine aquaculture systems that increasingly play a central role in emerging food supply challenges (Gentry et al., 2017).

Zebrafish are versatile research organisms that can be experimentally infected with viruses from heterologous fish and mammals (Palha et al., 2013; Phelan et al., 2005; Sanders et al., 2003). Larvae are mostly transparent, allowing for the visualization of antiviral immune responses in live transgenic animals (Palha et al., 2013). Two naturally occurring viruses have been detected in diseased adult zebrafish from a research facility and the pet trade (Bermudez et al., 2018; Binesh, 2013), and a third virus was recently discovered from an asymptomatic adult (Altan et al., 2019). These observations highlight the possibility that zebrafish regularly interact with viruses even in carefully maintained environments like research facilities, which might manifest in the induction of host antiviral signaling.

To advance the visualization of immune responses as a tool for virus discovery, we generated transgenic zebrafish that report on antiviral interferon signaling. We observed seemingly spontaneous induction of the reporter in zebrafish larvae during normal rearing in our research facility. With RNA-sequencing, we discovered a picornavirus in animals exhibiting antiviral immune responses that was distantly related to all other picornaviruses known at the time of discovery. Beyond our research facility, we found virus reads in RNA-sequencing datasets derived from zebrafish housed in research institutions around the world. Most notable were virus matches in datasets from a recently acquired wild zebrafish and striking levels of picornavirus from a clonal laboratory line (CG2), pointing to both natural and immunocompromised hosts. Finally, we begin to connect our analysis of virus burden with a previously unrecognized and substantial effect on host gene expression enriched for antiviral immune responses in otherwise apparently asymptomatic hosts.

## Results

### Spontaneous induction of interferon responses in zebrafish larvae

We generated transgenic zebrafish that report on interferon signaling to visualize antiviral immune responses in live animals. Interferons are signaling proteins that induce the expression of hundreds of genes collectively termed interferon-stimulated genes, the products of which provide the main antiviral immune defenses in vertebrates (Secombes and Zou, 2017). The promoter from zebrafish interferon-stimulated gene 15 (*isg15*) was cloned upstream of the coding sequence for GFP in a construct that also drives constitutive GFP expression in the heart, which was inserted into the genome by *Tol2* transposon-mediated transgenesis (Figure 1A). As in other vertebrates, the zebrafish *isg15* promoter contains regulatory transcription factor binding site motifs called interferon-stimulated response elements (Seppola et al., 2007). To test the interferon responsiveness of *isg15:GFP* animals we cloned the coding sequence for a type I interferon (*ifnϕ1*) into an expression plasmid (Figure 1B) and injected this construct into transgenic animals. Most untreated animals exhibited GFP expression that was restricted to the heart, which confirmed that they carried the transgenic insertion (Figure 1C). In contrast, animals treated with interferon expressed GFP broadly (Figure 1D and Video 1), demonstrating that the *isg15:GFP* transgenic animals provide a conspicuous readout for antiviral immune signaling.

**Figure 1.**
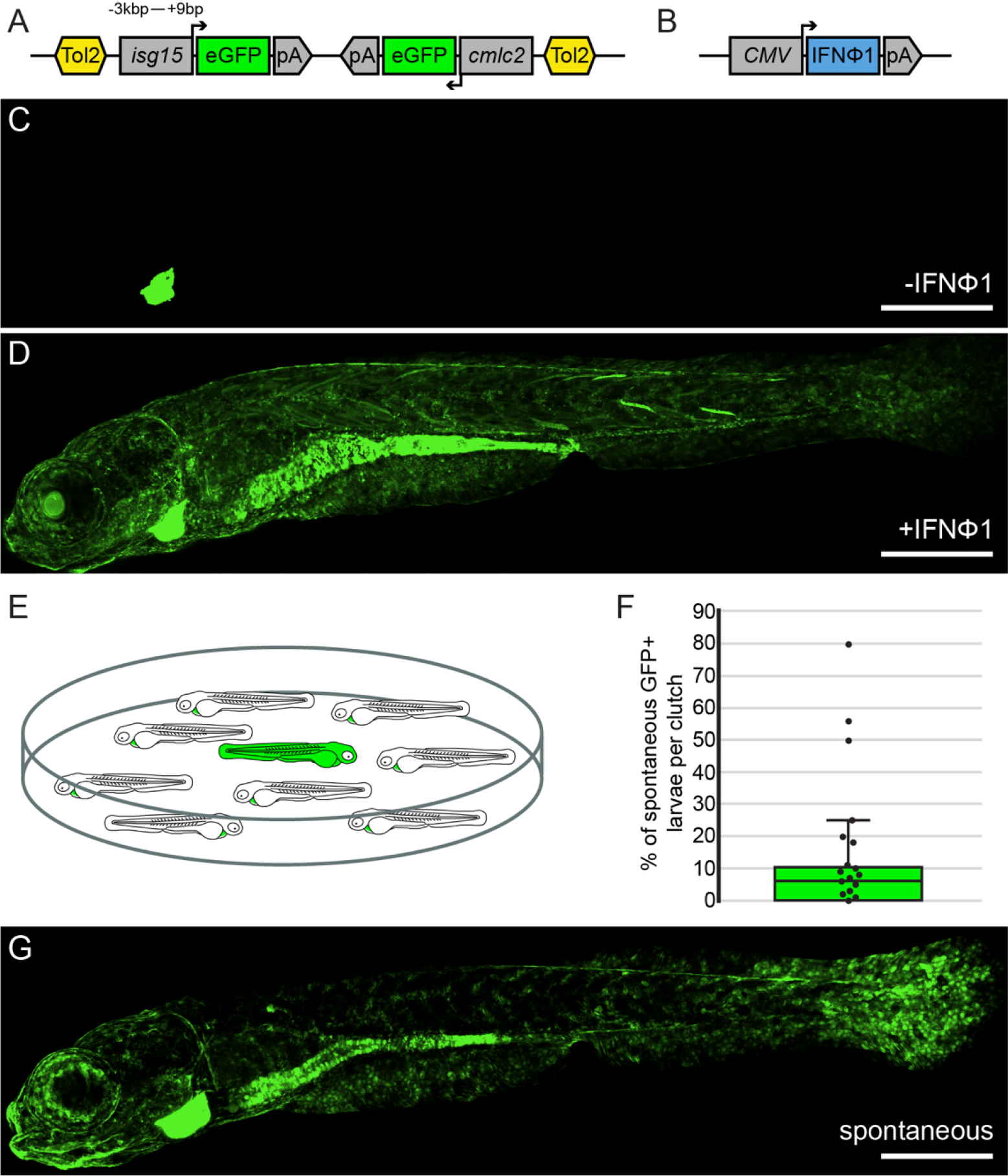
Induced and spontaneous GFP expression in transgenic zebrafish that report on interferon signaling. (A) Transgenic animals were generated with a Tol2 transposon construct bearing the promoter from *isg15* fused to the coding sequence for GFP along with a heart-specific (*cmlc2*) transgenesis marker in the opposite orientation. (B) Illustration of the construct used to drive type I interferon expression in zebrafish, with IFNϕ1 driven by the *CMV* promoter. (C) An untreated *isg15:GFP* zebrafish at 8dpf. (D) An IFNϕ1-treated *isg15:GFP* zebrafish at 8dpf. (E) Illustration of a dish of untreated *isg15:GFP* zebrafish larvae in which spontaneous GFP expression was observed. (F) Frequency at which spontaneous GFP was observed across independent clutches of *isg15:GFP* zebrafish. Each dot represents the percent of larvae within a clutch that spontaneously expressed GFP. (G) An untreated *isg15:GFP* zebrafish that spontaneously expressed GFP at 8dpf. All microscope images are maximum intensity projections of confocal Z-stacks. Scale bars are 500μm, dpf: days post fertilization.

We were surprised to observe that some untreated *isg15:GFP* transgenic fish spontaneously expressed GFP in multiple tissues by eight days post-fertilization (8dpf), which typically occurred at low frequency among animals from clutches in shared environments (Figure 1E). The distribution of spontaneous expression across multiple independent clutches was skewed towards low frequencies but varied widely (Figure 1F). Spontaneous GFP expression in *isg15:GFP* animals was most evident in tissues exposed to the environment, particularly the intestine and other epithelial surfaces (Figure 1G).

### Discovery of a picornavirus from zebrafish exhibiting antiviral immune responses

We hypothesized that the spontaneous GFP expression we observed in *isg15:GFP* animals reflected instances of virus infection. To identify prospective viruses, we isolated RNA from GFP-positive and GFP-negative *isg15:GFP* and wild type (WT) animals for RNA-sequencing (RNA-seq), generating approximately 175 million reads. We assembled reads that did not map to the zebrafish genome into approximately 750,000 larger contigs and assigned a preliminary species identity for each contig by taking the best hit from nucleotide or amino acid sequence similarity searches. The majority of these contigs (95%) were most similar to zebrafish sequences (Figure 2 - figure supplement 1), which evidently failed to map to the zebrafish genome as individual reads during alignment. Contigs with similarity to bacteria sequences were mostly from genera that are known constituents of the zebrafish microbiome (Roeselers et al., 2011). One contig 1,182bp in length had 33% amino acid identity (e-value 5×10^-40^) to the helicase of rosavirus 2, a picornavirus recovered from humans (Lim et al., 2014). No other contigs had significant similarity to known exogenous viruses.

We used the contig sequence with similarity to rosavirus 2 to design rapid amplification of cDNA ends (RACE) experiments that we carried out with cDNA template from spontaneously GFP-positive *isg15:GFP* animals. We used the resulting sequences to assemble a non-segmented positive-sense RNA genome of approximately 8kbp that encodes a polyprotein with features that are characteristic of picornaviruses (Figure 2A). Sequences in the 5’ UTR showed evidence for strong local secondary RNA structures. An IRES was predicted within the 5’ UTR, although no sequence similarity between the 5’ UTR and the IRES elements of other picornaviruses was observed. The structural and some nonstructural genes have sequence similarity to other picornaviruses, but portions of the genome do not share significant nucleotide or amino acid sequence similarity with anything and lack recognizable protein domains. These observations indicate that the virus we identified from *isg15:GFP* animals is distantly related to known picornaviruses.

**Figure 2.**
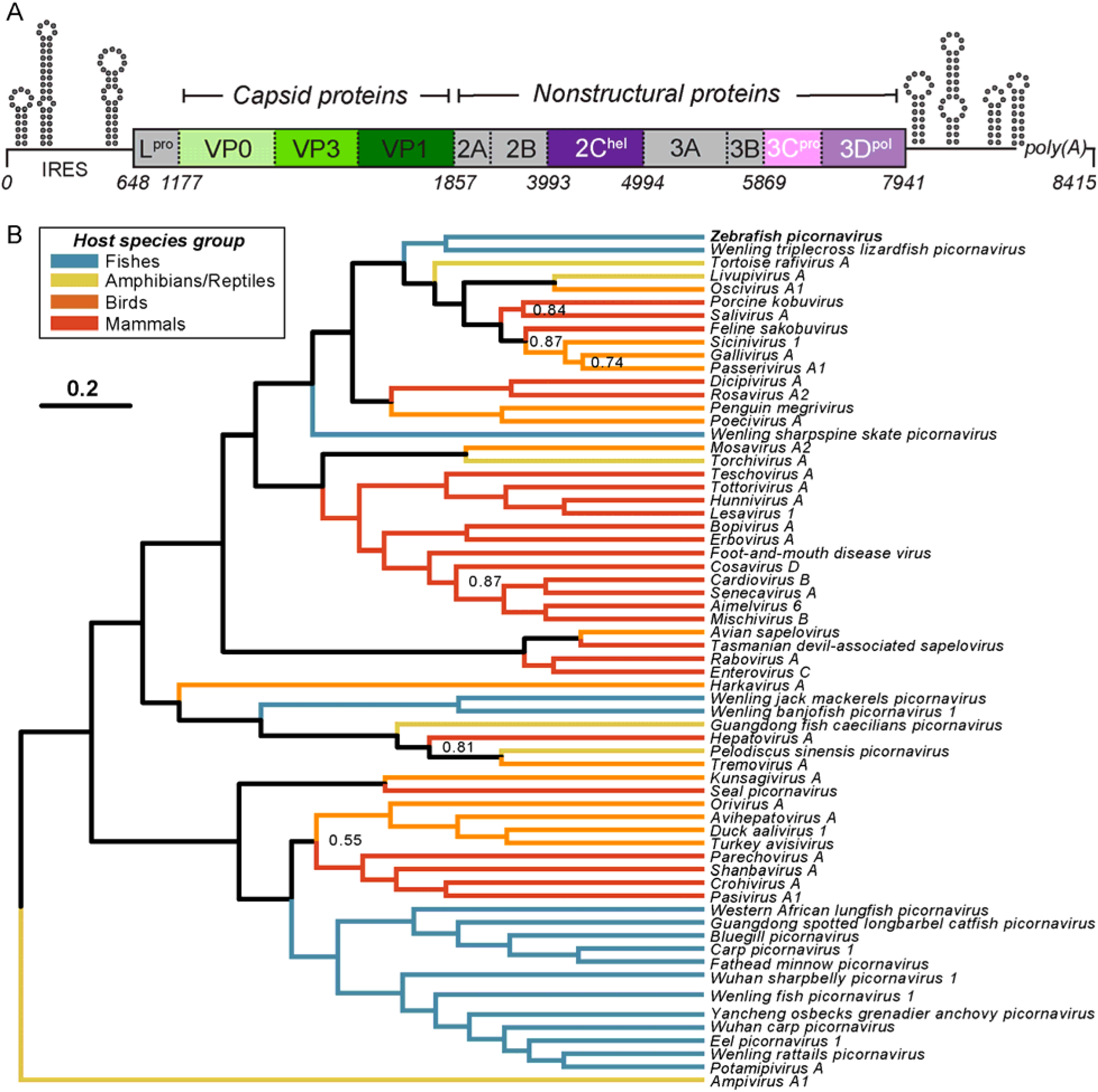
A new picornavirus sequenced from zebrafish larvae exhibiting antiviral immune responses. (A) Genome organization of *Zebrafish picornavirus*. Untranslated regions of the genome are depicted as lines. Schematics above lines represent secondary structures in the UTRs with high base-pairing probabilities based on minimum free energy predictions. The 5’ UTR is predicted to contain an internal ribosome entry site (IRES). Coding regions are depicted as boxes. Dashed lines indicate approximate cleavage positions within the polyprotein. Green and purple colors denote regions that have homology to other picornaviruses. Grey regions lack recognizable protein domain structures and do not share sequence similarity with any other organism. Names in boxes are based on similarity to other picornavirus proteins or are inferred based on the conserved nature of picornavirus genome structures. Numbers below the boxes refer to nucleotide positions. (B) Maximum clade credibility tree of picornavirus 3C and 3D amino acid sequences. Branches are colored based on the group of vertebrates that serve as hosts. Most nodes were strongly supported, with posterior probabilities >0.90 (values not indicated in the tree for clarity). Nodes with weaker support, (<0.90) are highlighted in the tree with their estimated support. Scale is in expected number of substitutions per site.

Subsequent to our sequence analyses described above, a contemporaneous study published a picornavirus sequence that was obtained through metagenomic sequencing of enriched viral capsid-protected nucleic acids from the intestine of an asymptomatic adult zebrafish (Altan et al., 2019). The authors of the study called this virus strain Zebrafish picornavirus (ZfPV), which is 97% identical to the nucleotide sequence of the virus that we assembled from *isg15:GFP* larvae. We describe the picornavirus in the present study as ZfPV for the sake of consistency.

To assess the evolutionary relationship of ZfPV to other members of the *Picornaviridae* family, we performed phylogenetic analyses including at least one species from all ratified picornavirus genera (Zell et al., 2017) and all available sequences from picornaviruses that have been isolated from fish (Figure 2B). Bayesian and maximum likelihood phylogenies were largely congruent (Figure 2 - figure supplement 2). We identified a large clade of viruses that appears to be fish-specific, which does not include ZfPV. ZfPV is part of a clade of viruses that infect a mixture of other vertebrate groups. The examples of incongruities between viruses and host groups in this phylogeny are consistent with patterns of virus host switching, which was previously estimated to occur frequently in picornavirus evolution (Geoghegan et al., 2017). The Wenling triplecross lizardfish picornavirus is the sister species to ZfPV in this phylogeny, although they only share 36% amino acid identity in a pairwise alignment of 3CD and are therefore only distant relatives (Figure 2 - figure supplement 3). Overall, these observations demonstrate that ZfPV is highly diverged from the other picornaviruses that are currently known.

### Widespread prevalence of ZfPV in publicly available zebrafish RNA-seq datasets

Our observations of spontaneous induction of GFP expression in independent clutches of *isg15:GFP* larvae were made separately over the course of several months (Figure 1F), which indicates that ZfPV is endemic to our research facility. We surveyed the prevalence of ZfPV in other research facilities by searching for ZfPV reads in zebrafish RNA-seq datasets in the Sequence Read Archive (SRA). Our search revealed widespread prevalence of ZfPV in research facilities around the world (Figure 3A). ZfPV reads were identified in RNA-seq datasets derived from a variety of zebrafish strains that were housed in various locations. We were particularly interested by the identification of ZfPV reads in a dataset (SRR891512) from a zebrafish line called Assam that appears to have been recently caught from its natural habitat in Northeastern India or was recently derived from the wild (Patowary et al., 2013; Sharma et al., 2019). The identification of ZfPV reads from a zebrafish that was either recently caught or derived from the wild supports the possibility that ZfPV also infects zebrafish in natural habitats.

**Figure 3.**
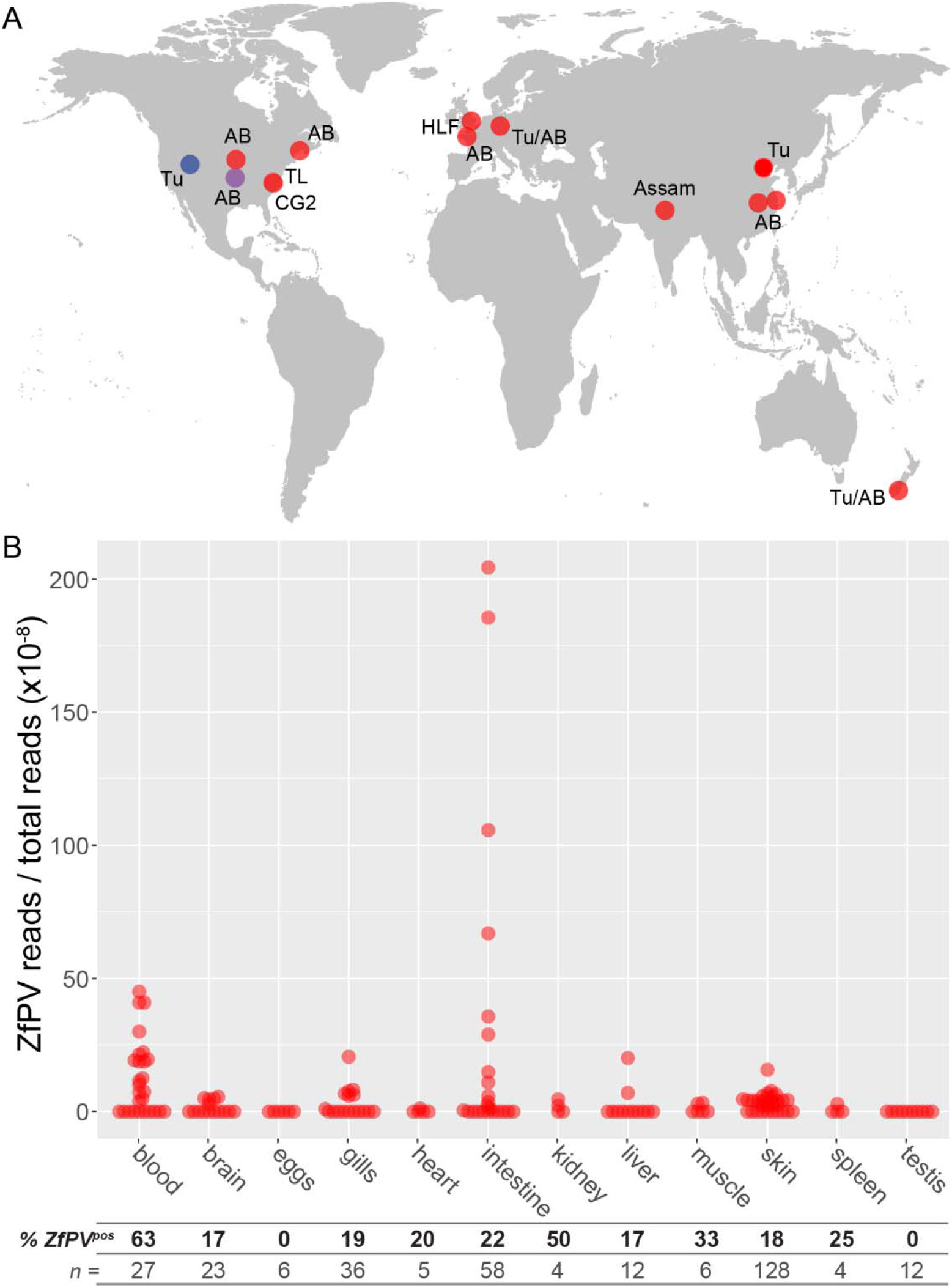
ZfPV is widespread in publicly available RNA-seq datasets. (A) Locations of institutions that have submitted zebrafish RNA-seq projects to the Sequence Read Archive (SRA) that contain reads from ZfPV. Red dots denote locations in which ZfPV reads were identified in the SRA. The blue dot denotes the location of the present study where ZfPV was initially discovered. The purple dot denotes the location of a contemporaneous study where ZfPV was also discovered (Altan et al., 2019). Names next to the dots refer to the strain of zebrafish that was sequenced. (B) Number of ZfPV reads normalized by library size that were identified in the SRA from adult tissues. Each dot represents a single sequencing sample. The percentage of SRA samples that contained ZfPV reads and the total number of samples that were searched for each tissue are shown beneath the plot.

The widespread prevalence of ZfPV in zebrafish datasets indicates that zebrafish are commonly infected. To assess potential host range beyond zebrafish we searched for ZfPV in all available datasets in the SRA from fish intestinal tissues, which included 775 RNA-seq datasets from more than 100 species of fish and found no evidence for ZfPV in these hosts (Figure 3 - figure supplement 1), demonstrating that ZfPV is specific to sequencing datasets from zebrafish among all fish species currently represented in the SRA.

We identified ZfPV in RNA-seq datasets that were derived from several different zebrafish tissues, which allowed us to evaluate the tissue tropism of the virus. The intestine had the highest number of virus reads (Figure 3B), which is consistent with complementary *in situ* hybridization analyses (Altan et al., 2019) and indicates that ZfPV may predominate as an enteric virus. However, we also observed ZfPV reads in datasets from several additional tissues, including those exposed to the environment such as the gills and skin but also internal tissues including the brain. In contrast, no virus reads were identified in datasets from eggs or testis. These patterns suggest that ZfPV may disseminate from the intestine to a variety of other tissues.

### The CG2 zebrafish strain harbors higher levels of ZfPV than WT animals

Two SRA datasets from BioProject PRJNA280983 (Dirscherl and Yoder, 2015) stood out during our survey for ZfPV in that they had exceptionally high numbers of virus reads (Figure 4A). These datasets were derived from the clonal CG2 strain of zebrafish (Mizgirev and Revskoy, 2010), and are not plotted in Figure 3B because they were generated from a mixture of tissues. The CG2 strain is genetically distinct from Tubingen and other WT lines in that CG2 animals harbor an alternative Major Histocompatibility Complex (MHC) haplotype, which has previously been predicted to confer differences in immunity-related phenotypes (Dirscherl and Yoder, 2015; McConnell et al., 2016, 2014). When we aligned reads from the two CG2-derived datasets to the ZfPV genome we recovered the entire genomic sequence up to nearly 165X coverage (Figure 4A, and GenBank accession number TPA: BK011179).

**Figure 4.**
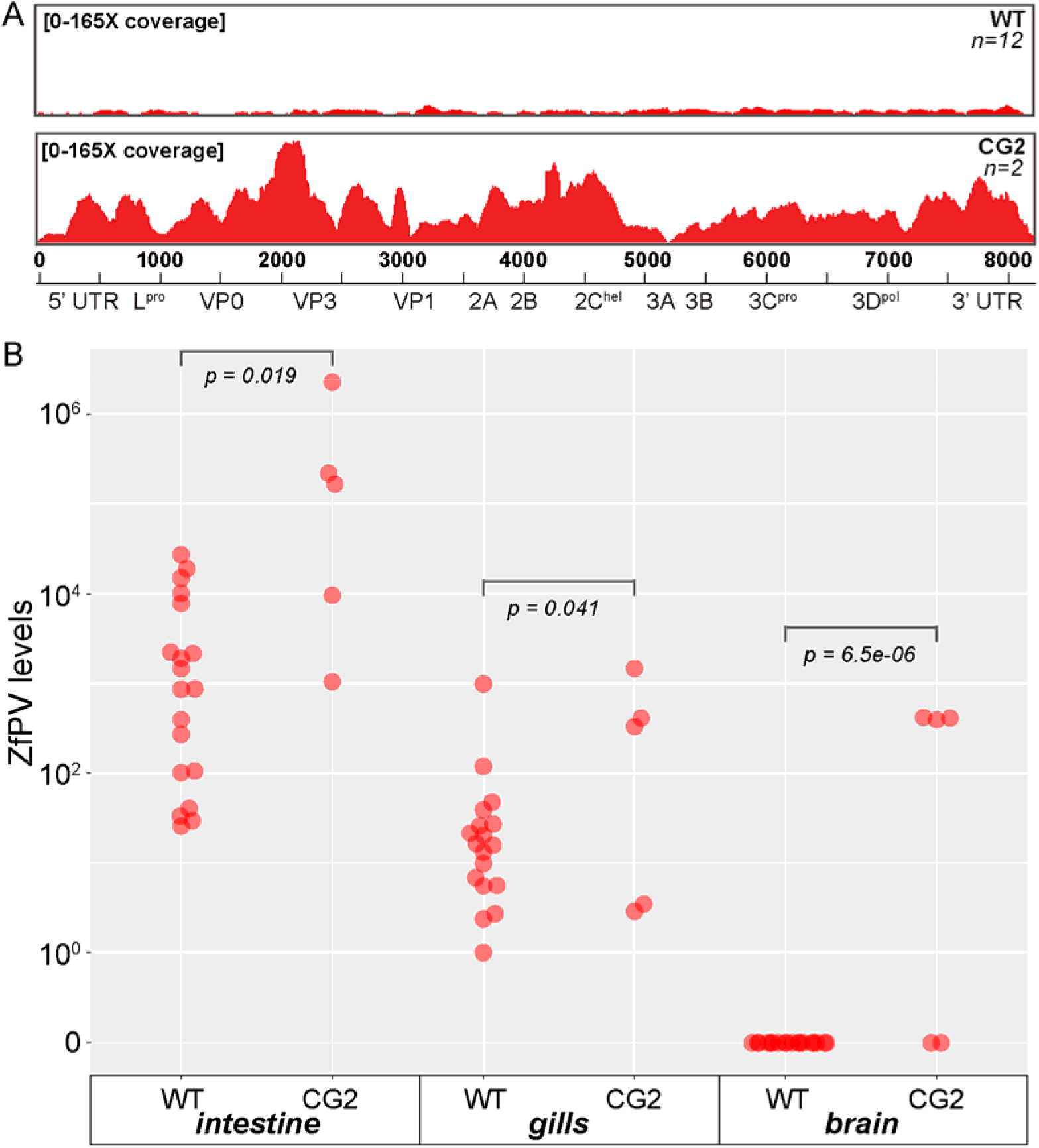
Virus levels are higher in the CG2 strain compared to WT zebrafish. (A) Coverage of the ZfPV genome by reads derived from all 12 WT intestine SRA datasets that had virus reads from Figure 3 compared to reads in the SRA derived from a single CG2 animal. Nucleotide positions and genomic features of ZfPV are shown below the coverage plots. Scale is in number of reads that cover each position. (B) Quantitative real-time PCR quantification of ZfPV levels in tissues dissected from adult WT and CG2 animals. Each dot represents the relative amount of virus in one individual. Virus levels are scaled by normalizing to the sample with the lowest non-zero amount of ZfPV. P-values are reported from unpaired t-tests assuming equal variance.

In comparison to the CG2 datasets, the majority of RNA-seq datasets that we queried in the SRA either had no ZfPV reads or very few reads relative to the total (Figure 3B). This is perhaps not surprising given that all of these datasets were generated to investigate host gene expression, and that virus RNAs typically account for a miniscule fraction of the total RNAs extracted from vertebrates for metagenomic sequencing (Geoghegan et al., 2018; Shi et al., 2018). To illustrate the relative rarity of virus reads even in the tissue with the highest ZfPV signal, we combined all 12 intestine-derived datasets that contained ZfPV reads from WT animals (Tubingen strain) and aligned these to the ZfPV genome. In contrast to the two CG2 datasets, aligning all 12 ZfPV-positive WT intestine datasets yielded low coverage across the ZfPV genome, with some regions that lacked coverage altogether (Figure 4A).

The higher degree of ZfPV coverage we observed for reads from CG2 datasets compared to WT datasets led us to hypothesize that the CG2 strain might harbor higher levels of virus than WT animals. To test this hypothesis, we isolated intestines, gills, and brains from adult CG2 and WT animals, extracted RNA, and generated cDNA to measure virus levels by quantitative real-time PCR. We detected ZfPV in the intestines and gills of every animal included in these experiments, reinforcing the seemingly ubiquitous nature of this virus in research facilities (Figure 4B). In support of our hypothesis, we observed higher levels of ZfPV in CG2 for all three tissues surveyed, including the brain where virus was only detected in CG2 animals. These data indicate that the large number of ZfPV reads that we observed in RNA-seq datasets from a CG2 animal appears to reflect a trend for the clonal and potentially immunocompromised strain, highlighting host genetic variation influencing virus levels.

### ZfPV infection alters gene expression in *isg15:GFP* and SRA RNA-seq datasets

We discovered that *isg15:GFP* animals were infected through observing induction of GFP expression, which established that ZfPV elicits a host immune response. To comprehensively quantify the effects that ZfPV infection has on host gene expression, we compared gene expression levels in spontaneously GFP-positive *isg15:GFP* animals to *isg15:GFP* and WT animals from the same clutches that were GFP-negative. From the same RNA-seq reads that we used to discover ZfPV, we identified more than 500 host genes (at an adjusted p-value of less than 0.05) induced by at least two-fold in virus-infected *isg15:GFP* animals compared to uninfected animals (Figure 5A). The most significant functional classes that were enriched among genes induced by infection were related to interferon signaling and defensive responses against viruses (Figure 5 - figure supplement 1). These results reveal that beyond the induction of *isg15:GFP*, naturally occurring picornavirus infections elicit extensive antiviral transcriptional responses in zebrafish larvae.

**Figure 5.**
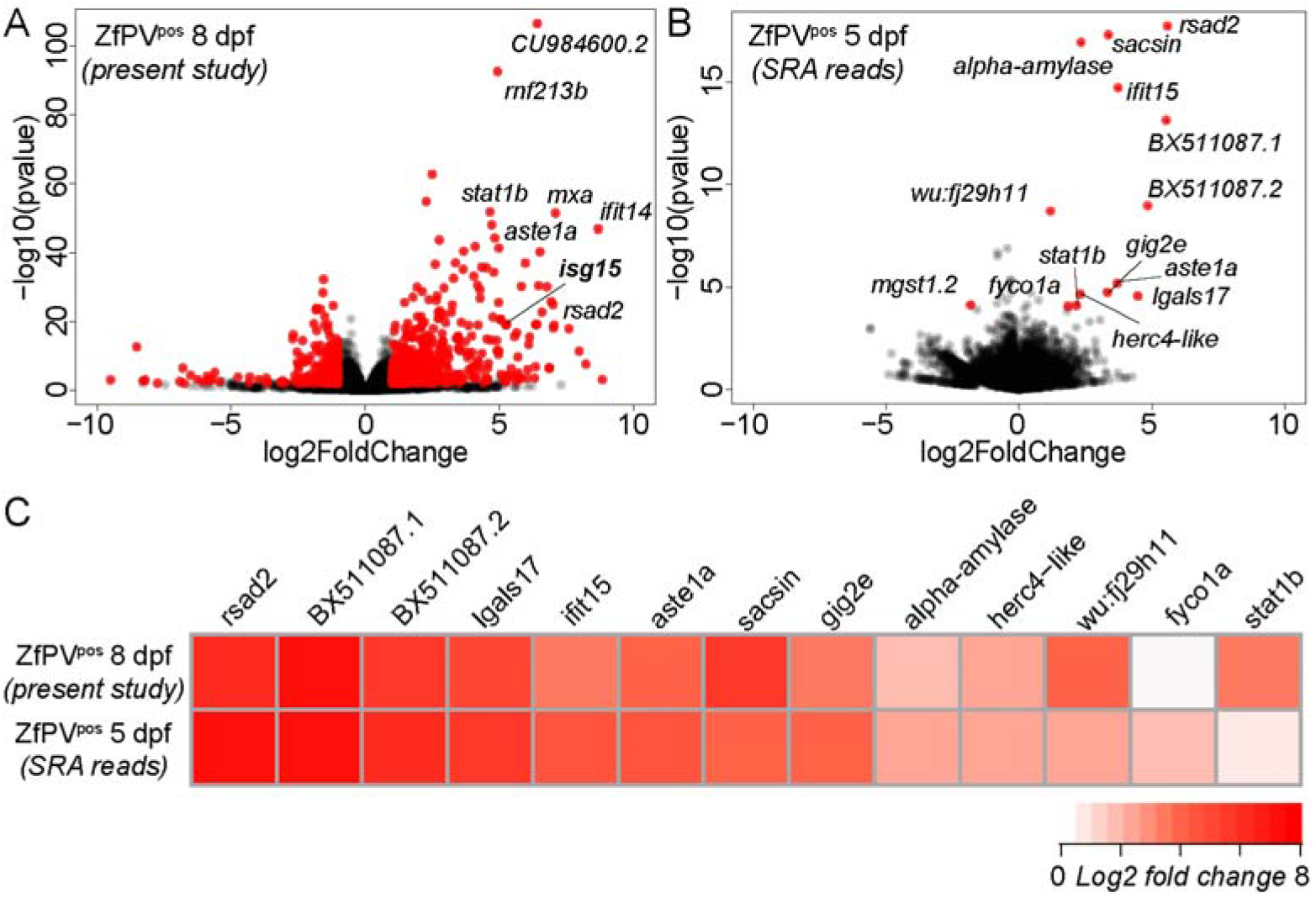
ZfPV infection elicits interferon responses in *isg15:GFP* animals and within replicates in SRA datasets. (A) Differential gene expression in *isg15:GFP* animals spontaneously expressing GFP compared to GFP-negative animals 8dpf. Only GFP-positive samples contained ZfPV reads. (B) Differential gene expression in 5dpf replicates that contained ZfPV reads compared to replicates that did not from BioProject PRJEB12982. Genes that were differentially expressed greater than 2-fold with adjusted p-values less than 0.05 are plotted in red. (C) Comparison of differential gene expression between 8dpf and 5dpf animals for all genes induced by ZfPV in 5dpf animals.

To extend our analysis of ZfPV-induced genes to previously published RNA-seq datasets from fish not previously known to contain virus, we searched for datasets with biological replicates that vary in the presence of ZfPV. We performed differential gene expression analyses with data from a developmental time course (White et al., 2017) by comparing multiple 5dpf replicates that contained ZfPV reads to replicates that did not, and identified a set of genes that were significantly induced in the replicates positive for ZfPV (Figure 5B). Of the 13 genes significantly induced by ZfPV in 5dpf larvae, 12 were also induced in ZfPV-positive 8dpf *isg15:GFP* larvae (Figure 5C). Furthermore, none of these genes were significantly induced in 5dpf larvae that were infected with the pathogenic bacteria *Mycobacterium marinum* (Figure 5 - figure supplement 2), and as such constitute a specific response to virus infection. Together, these observations reveal a signature of ZfPV infection in RNA-seq datasets comprised of virus sequences and transcriptional changes in the host.

## Discussion

Several metagenomic surveys revealed that members of *Picornaviridae* infect diverse fish and other vertebrate species (Geoghegan et al., 2018; Shi et al., 2018; Zell, 2018), but experimental studies of picornaviruses are so far restricted to mammals. In combination with the wealth of tools developed in a productive history of experimental work with mammalian picornaviruses, our discovery in zebrafish provides a new system for investigating the biology of picornaviruses in a non-mammalian vertebrate host.

By carrying out virus discovery in a widely used research organism, we were able to extend our discovery of ZfPV in zebrafish larvae from our research facility to locations around the world. Our observation of virus reads in a zebrafish from India supports the idea that ZfPV circulates in the wild. This provides the opportunity to study natural virus-zebrafish interactions and build on previous *in vivo* studies of zebrafish infections by several viruses isolated from other organisms (Van Dycke et al., 2019; Varela et al., 2017). While definitive classification of ZfPV as a virus that infects wild zebrafish awaits direct sampling of animals in natural habitats, the widespread prevalence of ZfPV in research facilities alone offers the possibility of studying host-virus evolution in a variety of controlled environments.

One of the notable findings from our study was the exceptional number of virus reads in datasets from the CG2 strain of zebrafish (Figure 4A). This observation led us to survey ZfPV levels in tissues that we extracted from several CG2 and WT adult animals, which revealed that ZfPV levels are regularly higher in the CG2 strain than in WT animals (Figure 4B). WT and CG2 animals are genetically distinct in at least one important way: WT animals carry a heterogenous mix of distinct MHC core haplotypes and CG2 animals are homozygous for a single core MHC haplotype (McConnell et al., 2014). Variation at the MHC locus is known to alter susceptibility to infectious diseases in mammals, birds, and fish (Grimholt et al., 2003; Hunt et al., 2010; Matzaraki et al., 2017), and zebrafish genomes contain extensive MHC gene diversity (McConnell et al., 2016). Thus, our observation that the CG2 strain tends to harbor higher levels of ZfPV than WT animals expands on our discovery of a new virus by revealing candidate host factors that influence susceptibility to infection.

ZfPV appears to infect primarily as an enteric pathogen of zebrafish, but the full range of tissue tropism is complex (Figure 3B). This is common for picornaviruses as exemplified by poliovirus, which is thought to primarily cause gastrointestinal infections in humans but can then spread to the blood and occasionally other tissues including the central nervous system in a manner that is influenced by the host immune system and other factors (Racaniello, 2006). Our observations are consistent with this pattern for ZfPV, which we detected at the highest levels in the intestine but also in several other tissues including the brain (Figure 3B). Intriguingly, we detected ZfPV in brain tissue that we collected from CG2 fish but not from WT fish (Figure 4C). We interpret this expanded tissue tropism as linking the susceptibility of the CG2 strain to a dynamic infection process that involves both virus and host factors.

The ZfPV genome encodes several predicted proteins that lack recognizable sequence similarity to any known protein or functional domain. Picornaviruses are known to inhibit host interferon responses in mammals in part by directly cleaving ISG15 (Swatek et al., 2018), and we therefore suspect that ZfPV proteins interact with specific members of the teleost fish interferon system. In addition to yielding insights into the functions of picornavirus proteins in this system, investigating the non-coding regions of the genome could provide new molecular tools for understanding the functions of proteins. An internal ribosome entry site (IRES) and a 2A “self-cleaving” peptide from mammalian picornaviruses are both widely used tools in molecular biology (Parks et al., 1986; Ryan et al., 1991), and analogous functions with distinct applications might be encoded the ZfPV genome.

Many of the strengths of the zebrafish model system for studying host-pathogen interactions lie in the transparency and small size of larval animals (Levraud et al., 2014; Tobin et al., 2012), which is the life stage from which we discovered ZfPV. In a study contemporaneous with ours, Altan and colleagues identified a virus in zebrafish gut tissue in a metagenomic investigation of an asymptomatic adult individual (Altan et al., 2019). The genomic sequence for this virus exhibits 97% nucleotide identity with the picornavirus we discovered and therefore appears to be the same species. We observed ZfPV infections in both adult and larval animals, opening up multiple levels of experimental opportunities for studying the dynamics of infection. Furthermore, our work is unique in revealing both virus and its effects on host biology.

This study highlights a set of ZfPV-induced host genes that encode for proteins with a clear signature of interferon- and virus response-related functions. The possibility that unrecognized infections by naturally occurring viruses might cause unexplained variation in experiments with zebrafish has been a longstanding concern (Crim and Riley, 2012). We observed gene expression differences within experimental replicates in previously published RNA-seq datasets likely caused by the presence of ZfPV (Figure 5B), which supports the idea that unrecognized virus infections contribute to experimental variation in zebrafish studies. The full range of effects that ZfPV might have in zebrafish gene expression studies is unclear, as the majority of RNA-seq experiments in the SRA do not include enough replicates with and without virus reads to quantify significant differences. However, monitoring the presence of ZfPV in future zebrafish gene expression studies to account for host responses to cryptic infections should now be straightforward.

ZfPV is likely one of many viruses circulating in zebrafish populations, and *isg15:GFP* fish could be useful sentinels for discovering additional viruses that infect zebrafish from a variety of environments. Variations on this strategy could lend further sensitivity and versatility to the discovery process, such as through the combinatorial use of promoters that report on distinct aspects of infection or by generating reporters in immunocompromised host backgrounds. More generally, our study outlines an adaptable strategy that could be deployed in many organisms to illuminate the ubiquitous but typically invisible engagements between viruses and hosts.

## Materials and methods

### Fish maintenance and transgenesis

Zebrafish (*Danio rerio*) were maintained in accordance with approved institutional protocols under the supervision of the Institutional Animal Care and Use Committee (IACUC) of the University of Utah, which is fully accredited by the AAALAC. Wildtype zebrafish were from the Tubingen (Tu) strain. Embryos were collected following natural spawning, maintained at 28.5°C in E3 embryo medium (Westerfield, 2007). CG2 zebrafish (Mizgirev and Revskoy, 2010) were a gift from David Langenau and were euthanized upon arrival at the University of Utah from the Massachusetts General Hospital. *Tg(isg15:GFP)* zebrafish were generated with the Tol2kit as previously described (Kwan et al., 2007). A fragment containing 3.4kbp upstream of the start codon of *isg15* along with the first 9bp of coding sequence was amplified from Tu genomic DNA using primers *isg15_pro_F1 isg15_pro_R1* (Supplemental file 1). This fragment was cloned into a Gateway construct and fused in frame with GFP with the Tol2kit. 30pg of the *isg15:GFP* DNA construct was injected in Tu embryos at the onecell stage along with 25pg of transposase RNA. Three stable independent lines were generated and are collectively referred to as *isg15:GFP* animals in the text.

### Microscopy

Transgenic animals were identified and maintained by screening for GFP expression in the heart using an Olympus SZX12 stereo microscope outfitted with a 100W mercury bulb fluorescent light source (Olympus). The same microscope was used to observe induced and spontaneous GFP expression in *isg15:GFP* animals. For imaging, live 8dpf *isg15:GFP* animals were anaesthetized with tricaine and mounted in 1% low melt agarose. Animals were imaged using a Nikon A1 laser scan confocal microscope (Nikon Instruments) with an NA 0.45/10x objective. A 20μm step series to a depth of 160μm was acquired using a 48μm pinhole size at a resolution of 0.4 pixels per μm.

### Induction of type I interferon expression

The coding sequence of *ifnϕ1* (NM_207640.1), a type I interferon (Hamming et al., 2011), was synthesized and cloned downstream of a CMV promoter into the pCS2+ vector backbone. 50pg of plasmid was injected in *isg15:GFP* embryos at the single cell stage to induce interferon signaling. Animals were screened for GFP expression at the end of the first day and then daily up to 9dpf. Transgenic animals from the same clutches that were not injected were screened over the same period of time.

### Quantification of spontaneous GFP expression

Transgenic *isg15:GFP* embryos were collected following natural spawning and maintained at 28.5 °C in E3 embryo medium. Embryos were not bleached and hatched naturally. They were screened daily with a fluorescent stereo microscope for GFP expression outside of the heart (which expresses GFP constitutively) up to 9dpf. Animals that expressed GFP spontaneously were first observed at 8dpf. To quantify the frequency of spontaneous GFP expression animals were anaesthetized with tricaine on 8dpf or 9dpf. The number of GFP-positive and GFP-negative animals were counted for each clutch by manually inspecting each individual. There were approximately 200 individuals per clutch, and embryos were split across petri-dishes at the time of collection to contain 50 individuals per dish. Animals were euthanized after quantification of GFP expression.

### RNA sequencing

Four independent untreated clutches of *isg15:GFP* animals were screened for GFP expression and separated as GFP-negative and GFP-positive. 30 individuals were pooled per group. Three isg15:GFP and two WT Tubingen clutches were collected at 8dpf. RNA was extracted by mechanical lysis with TRIzol (ThermoFisher) and DNAse-treated. RNA quality was assessed by RNA ScreenTape assay (Agilent Technologies) yielding RIN^e^ scores of ≥ 9.6 for all samples. Libraries were prepared by the High Throughput Genomics Shared Resource at the University of Utah with the Illumina TruSeq Stranded Total RNA Ribo-Zero Gold kit, and were sequenced on an Illumina HiSeq 2500 by 125 cycle paired-end sequencing v4. Raw sequencing reads are available in the National Center for Biotechnology Informatics (NCBI) Sequence Read Archive (SRA) under BioProject accession number PRJNA575566 (in process).

### Virus genome assembly and annotation

FASTQ reads from all samples were mapped to the GRCz11 zebrafish genome assembly (Ensembl release 94) with STAR version 2.5.1 (Dobin et al., 2013). Unmapped reads were assembled into larger contigs with Trinity (Haas et al., 2013) with minimum contig length set to 150. All contigs were searched for nucleotide similarity to all sequences in the NCBI nucleotide (nt) database with BLASTn. For contigs that lacked similarity to known sequences at the nucleotide level, the longest ORFs and protein sequences were predicted for each contig with TransDecoder (https://github.com/TransDecoder/). Predicted proteins were then searched for similarity to sequences in the UniProt UniRef90 database with BLASTp. One contig (1.2 kbp in length) had significant but distant similarity only to a picornavirus protein. A full-length genome for this prospective virus was assembled by carrying out rapid amplification of cDNA ends (RACE) using cDNA prepared with the SMARTer 5’/3’ RACE kit (Takara Bio USA, Inc.) from the same RNA that was used for sequencing. The full-length genome was annotated with Genome Annotation Transfer Utility (Tcherepanov et al., 2006) and manually searched for conserved sequences and protein domains with BLAST and HMMER3 (Mistry et al., 2013). Minimum free energy structures in the UTRs were modeled with RNAfold in the Vienna RNA Websuite (Gruber et al., 2008). The internal ribosome entry site in the 5’ UTR was predicted with IRESfinder (Zhao et al., 2018). The full genome sequence for the virus we assembled from these sequencing data is available under GenBank accession number MN524064 (in process).

### Phylogenetic analyses

Phylogenies were estimated with the concatenated amino acid sequences from the picornavirus 3C protease and 3D RNA-dependent RNA polymerase. Sequences from at least one species of all known genera of picornaviruses were included, as well as all available sequences from picornaviruses that infect fish. All sequences were aligned with T-Coffee in PSI-Coffee homology extension mode (Di Tommaso et al., 2011), resulting in an alignment of 803 amino acid sites across 65 taxa. Evolutionary relationships were inferred using Bayesian phylogenetic analyses carried out in BEAST version 2.5.2 (Bouckaert et al., 2019). The Blosum62 substitution model was selected as the best-fit model for the alignment under the Akaike information criterion as determined by ProtTest version 3.4.2 (Darriba et al., 2011). The rate of evolution was set to 1.0 under a strict clock rate. A Yule model was implemented for the tree prior. The MCMC chain length was set to 10 million generations, storing every 5 thousand. Two independent runs were carried out from unique starting points. Convergence and mixing of phylogenetic tree topologies in these runs were assessed with the RWTY package (Warren et al., 2017) in R v3.6.0, and reached approximate effective sample sizes (ESS) of greater than 800. Convergence and mixing of all other parameters for these runs were determined with Tracer (Rambaut et al., 2018), and all reached approximate ESS values greater than 2000. A maximum clade credibility tree was generated with TreeAnnotator in the BEAST 2 package. Summarized and annotated trees were displayed with FigTree v1.4.4 (https://github.com/rambaut/figtree/releases).

### Searching the Sequence Read Archive for *Zebrafish picornavirus*

Publicly available RNA-seq datasets were searched with the NCBI SRA toolkit version 2.5.2 using sra-blast with the ZfPV genome as the query sequence. More than 1400 SRA datasets were queried in total, which were selected from the archive based on Boolean searches for the terms “zebrafish” and “intestine”, “gut”, “skin”, “gills”, “blood”, or “brain”. SRA datasets from other tissues were included if they were part of the same BioProject as those identified through the searches above. Non-zebrafish datasets were identified based on searching for “fish” and “intestine” or “gut”. 775 datasets from non-zebrafish fish species were investigated. Hits were considered significant if the e-value was ≥ 1e-10. The location of the institutions in which the zebrafish were housed in and the strains that were used for sequencing was determined by inspecting metadata associated with the SRA submissions and with descriptions in corresponding publications if available. To confirm that SRA dataset reads matched ZfPV, FASTQ files from SRA datasets with significant hits were downloaded and aligned to the ZfPV genome with Bowtie 2 (Langmead and Salzberg, 2012). SAM files were converted to BAM files and sorted with SAMtools (Li et al., 2009) to inspect manually with the Integrative Genomics Viewer (Robinson et al., 2011). We assembled a full zebrafish picornavirus genome sequence from BioProject PRJNA280983 raw reads, which is available under GenBank accession number TPA: BK011179.

### Quantification of virus levels by quantitative real-time PCR

Intestines, gills, and brains were dissected from WT (Tubingen strain) and CG2 adult animals (gift from David Langenau, MGH) of approximately one year of age. All animals were males raised in normal rearing conditions and did not exhibit any overt signs of disease. Tissues were homogenized in 250μl of PBS with a THb Tissue Homogenizer (Omni International) before adding 750μl of Trizol LS Reagent (ThermoFisher) to extract RNA. cDNA was prepared with the SuperScript IV kit (ThermoFisher) using 1μg of RNA for each sample. Quantitative real-time PCR was performed by amplifying a ZfPV target (a 161bp fragment in the 2C helicase) with primers ZfPV_F2 and ZfPV_R2 (Supplemental file 1) and a host reference gene target with primers elfa_F and elfa_R (McCurley and Callard, 2008) using Power SYBR Green Mastermix (ThermoFisher) on a QuantStudio 3 Real-Time PCR System (ThermoFisher). Primer efficiencies were measured by standard curve and were found to be 94% and 101% for elfa and ZfPV, respectively. The comparative C(T) method (Schmittgen and Livak, 2008) was used to determine relative levels of ZfPV per sample, which were normalized by dividing the values for all samples by the lowest non-zero value.

### Differential gene expression analyses

Transcript abundances were estimated from FASTQ files for all samples with Salmon version 0.14.0 (Patro et al., 2017) in mapping-based mode with the --validateMappings, --seqBias, and --gcBias options flagged. Reads were mapped to a transcript index built from the zebrafish transcripts annotated in Ensembl release 96 and modified to include the full genomic RNA sequence of ZfPV. Gene-level differential expression analysis was carried out by aggregating transcript-level quantification using DESeq2 version 1.24.0 (Love et al., 2014). Four *isg15:GFP* samples that spontaneously expressed GFP were compared to two *isg15:GFP* samples that did not express GFP and two non-transgenic WT Tu samples. Only the GFP-positive samples had reads that mapped to ZfPV. WT 5dpf RNA-seq datasets containing ZfPV reads were identified in BioProject PRJEB12982 (White et al., 2017) with sra-blast. FASTQ files for all 20 5dpf datasets from this BioProject were downloaded and differential gene expression analysis was carried out by comparing all replicates that contained ZfPV reads to those that did not.

## Acknowledgements

We thank Diego Mallo and Tyler Starr for input on phylogenetic analyses, Michelle Culbertson, Kristen Davenport, Diane Downhour, and Zoë Hilbert for critical feedback on the manuscript, Kristen Kwan and members of her lab for sharing Tol2 transgenesis reagents, and David Langenau and Alexandra Veloso for sharing the CG2 fish. We also thank the University of Utah Health Sciences Core Facilities for zebrafish husbandry and imaging support, and the Huntsman Cancer Institute High-Throughput Genomics Shared Resource for library preps and sequencing. This work was supported by grants awarded by the National Institutes of Health to KMB (5T32AI055434) and to NCE (R01GM114514). Additional support includes funding awarded to NCE by the Burroughs Wellcome Fund (1015462) and the HA and Edna Benning Presidential Endowed Chair.

## Figure Supplements

**Figure 2 - supplement 1.**
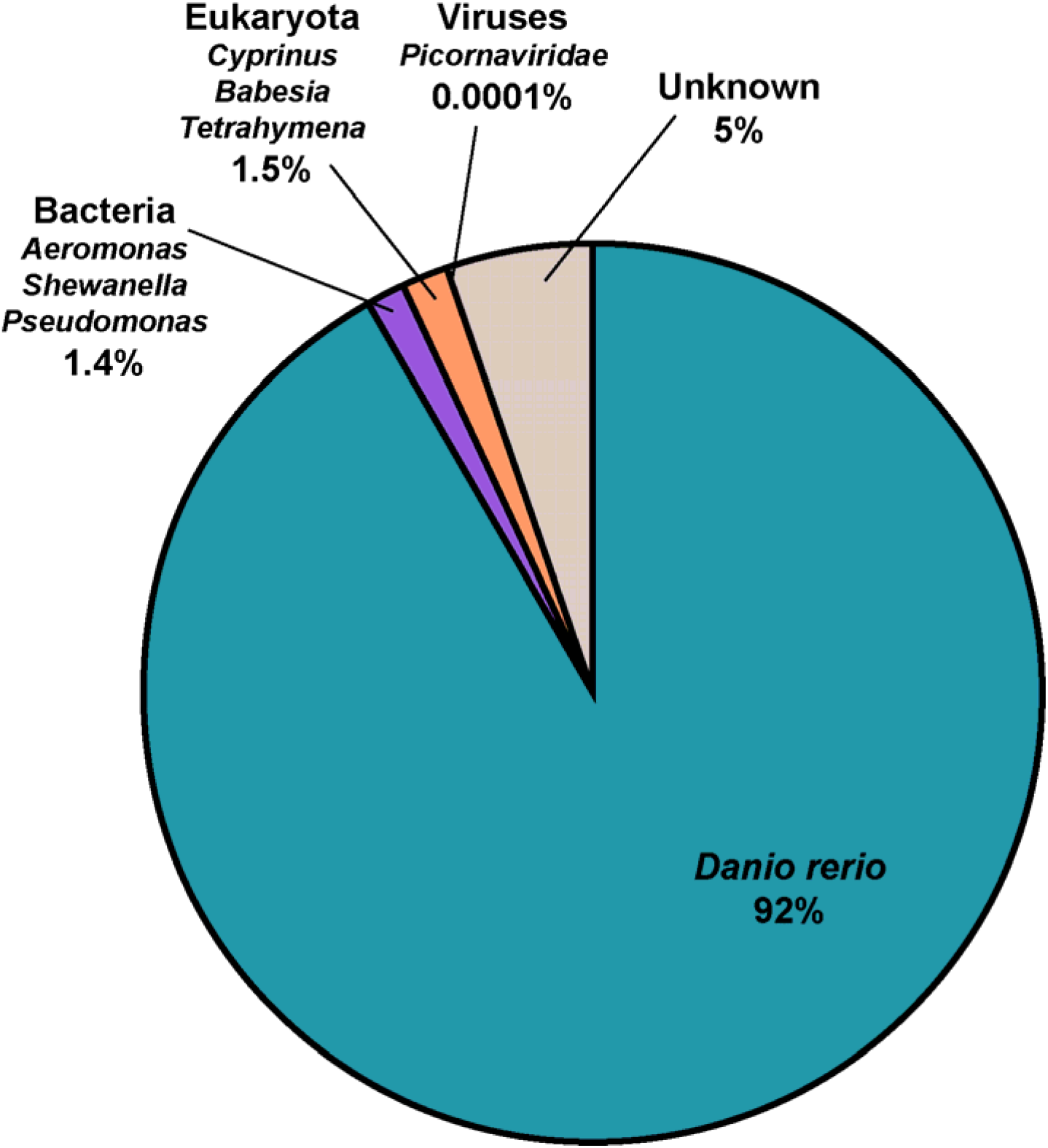
Species identities of all 737,608 contigs assembled from reads that did not map to the zebrafish genome during alignment. Most were determined to be from zebrafish and had failed to map as individual reads. The top 3 most abundant genera from other taxa are listed for Bacteria and Eukaryota. One contig was most similar to viruses. 5% of contigs had no similarity to any known nucleotide or amino acid sequence by BLAST. These were predominantly short (~150bp) and repetitive sequences.

**Figure 2 - supplement 2.**
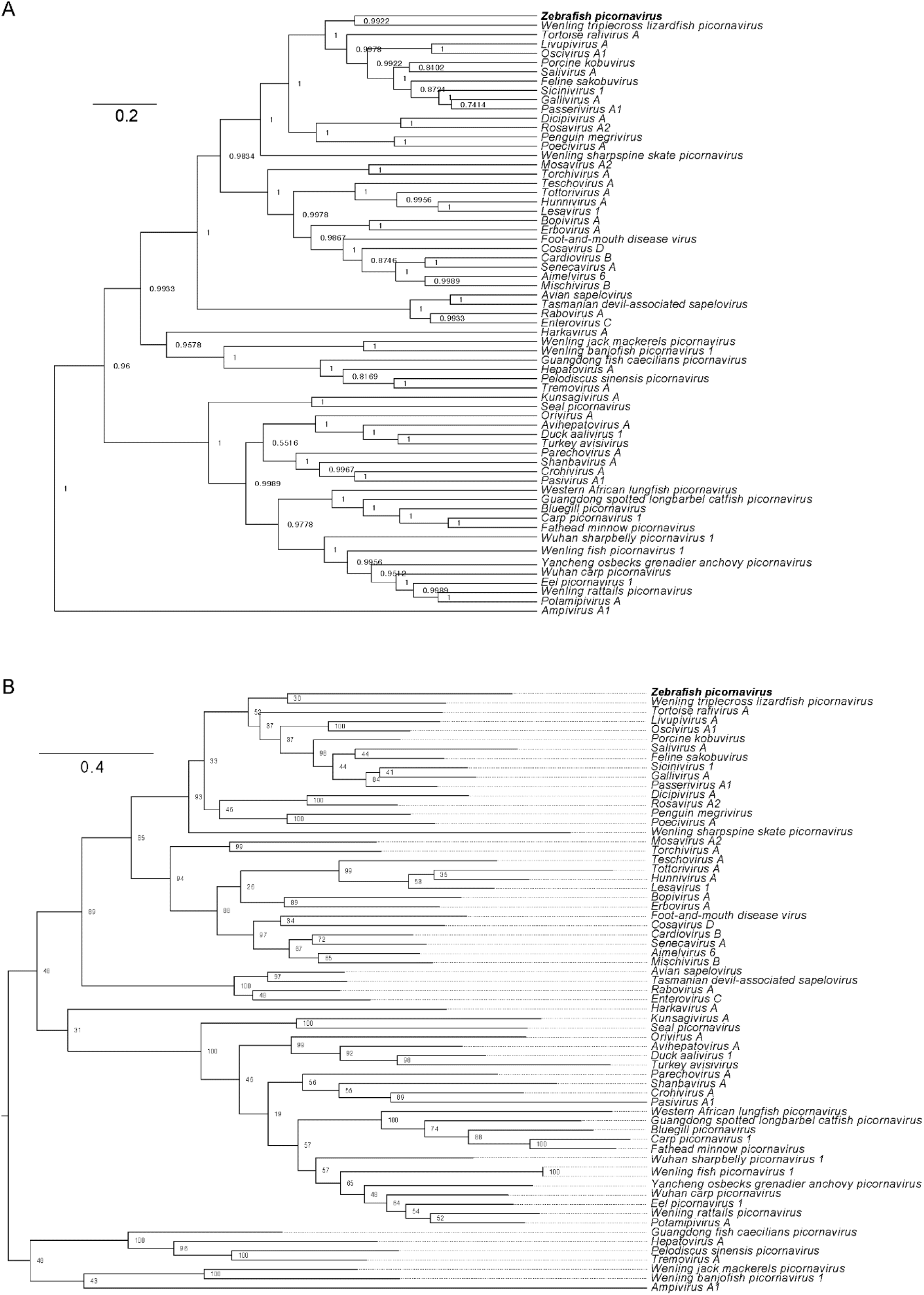
Bayesian and maximum likelihood phylogenies of picornaviruses. (A) The same maximum clade credibility tree shown in Figure 2B but fully annotated with posterior probabilities at all nodes. (B) Maximum likelihood phylogenetic tree of the same picornavirus 3C and 3D amino acid sequence alignment used for Bayesian inference. Phylogenetic relationships were estimated using the maximum likelihood method (with 100 bootstrap replicates) implemented in PhyML 3.0 (Guindon et al., 2010). Numbers at nodes are bootstrap values. The tree is midpoint rooted. Scale is in substitutions per site.

**Figure 2 - supplement 3.**
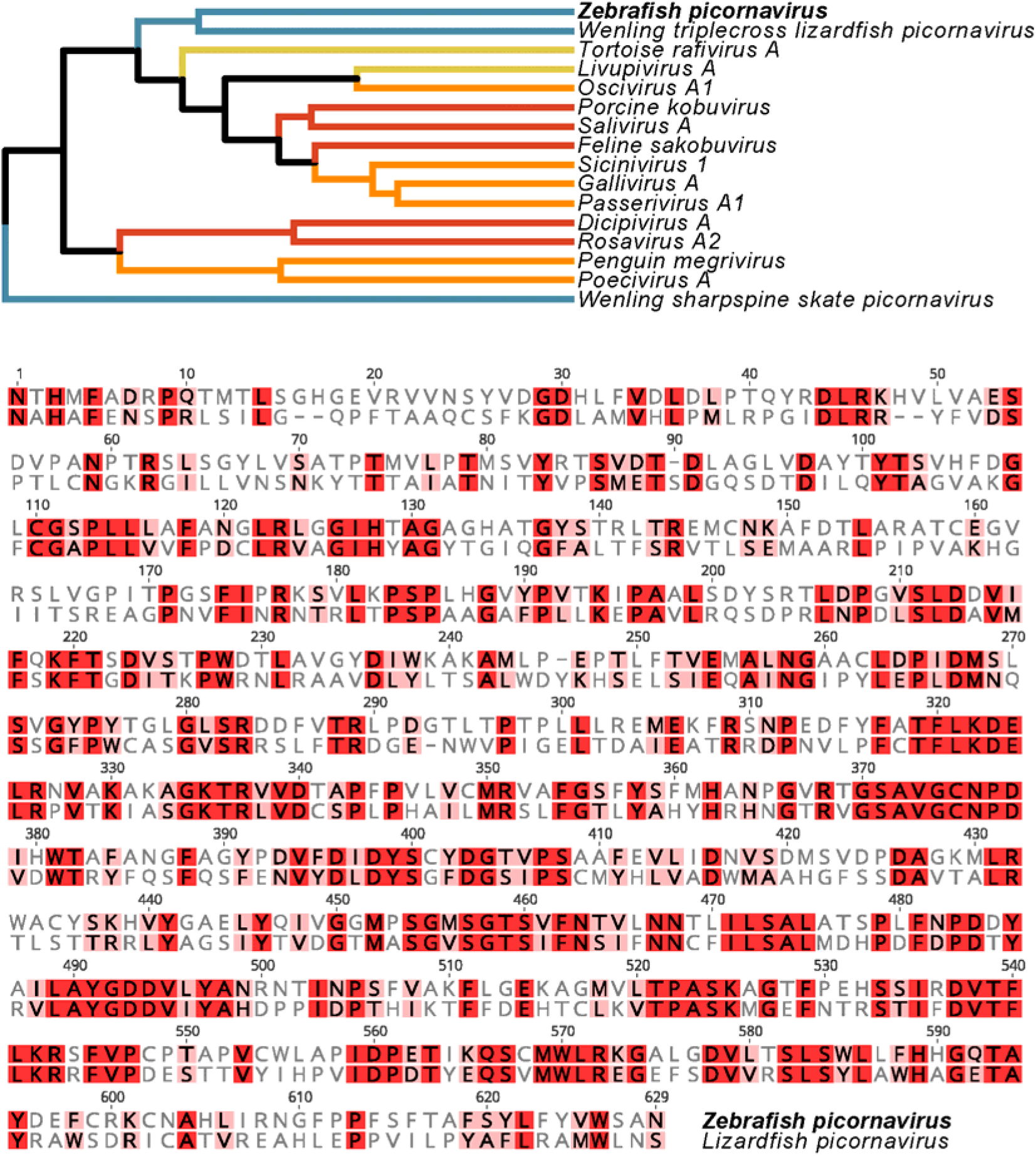
Pairwise alignment of the 3CD amino acid sequences from *Zebrafish picornavirus* and its closest relative, the *Wenling triplecross lizardfish picornavirus*.

**Figure 3 - supplement 1.**
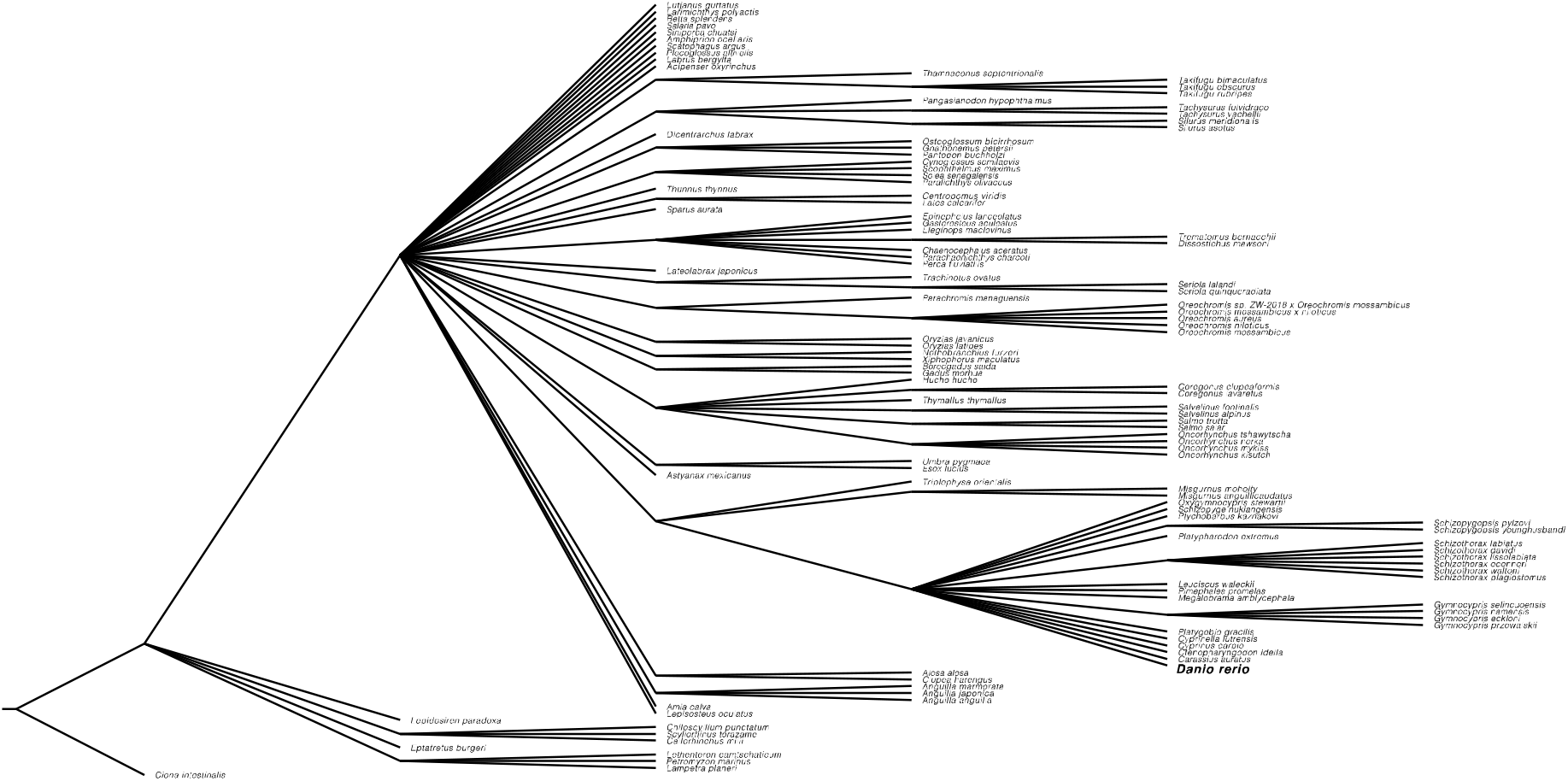
Common tree from the NCBI Taxonomy browser of all the fish species that were investigated for the presence of ZfPV. 775 non-zebrafish RNA-seq datasets were searched, but only zebrafish datasets contained ZfPV reads. The tree is rooted on *Ciona intestinalis* (which was not searched for ZfPV).

**Figure 5 - supplement 1.**
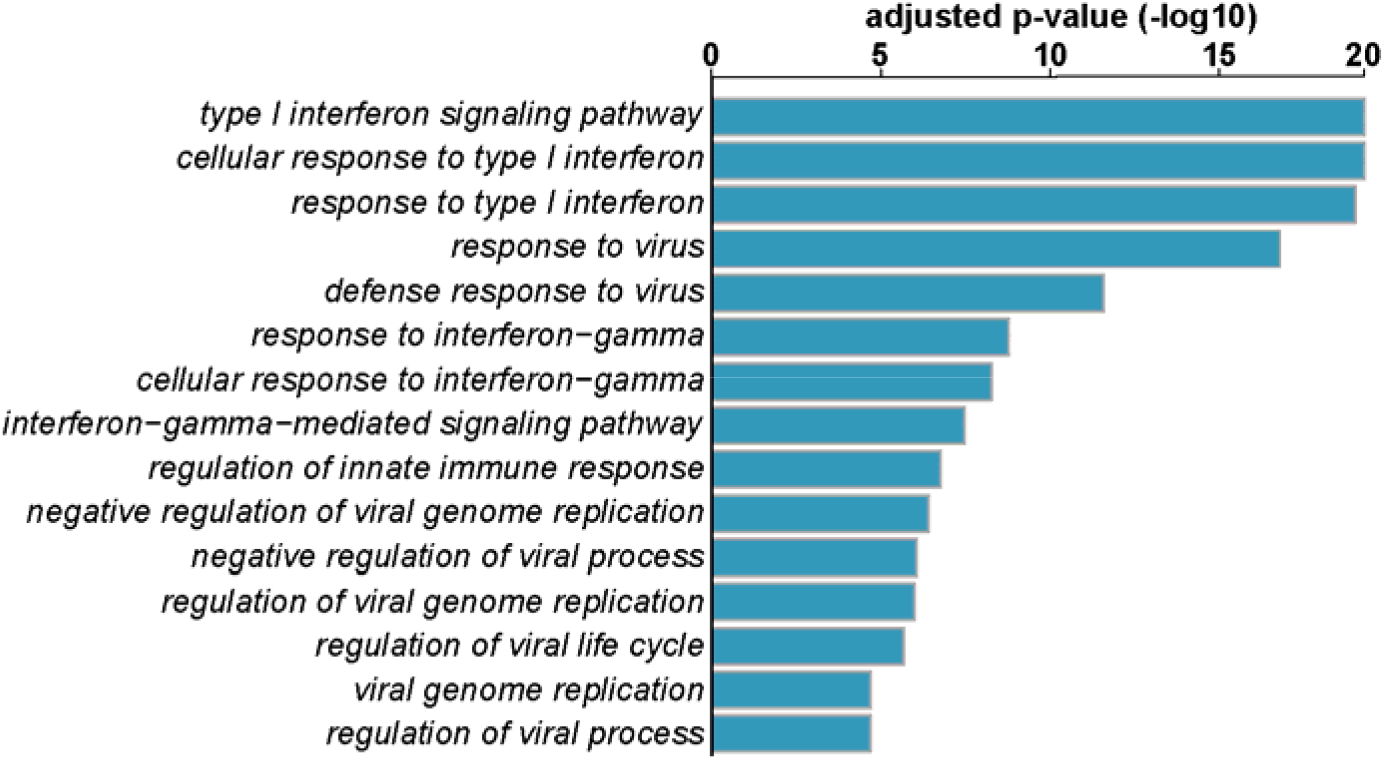
Gene set analysis of differentially expressed genes in *isg15:GFP* animals spontaneously expressing GFP compared to GFP-negative animals 8dpf. The top 15 significantly enriched Gene Ontology (GO) terms identified in the genes induced by ZfPV are plotted. The differentially expressed zebrafish genes were first compared to their homologs in human. GO terms were assigned based on functional classes having significant enrichment for both species. These analyses were carried out with the XGSA statistical framework for cross-species comparisons of gene sets (Djordjevic et al., 2016).

**Figure 5 - supplement 2.**
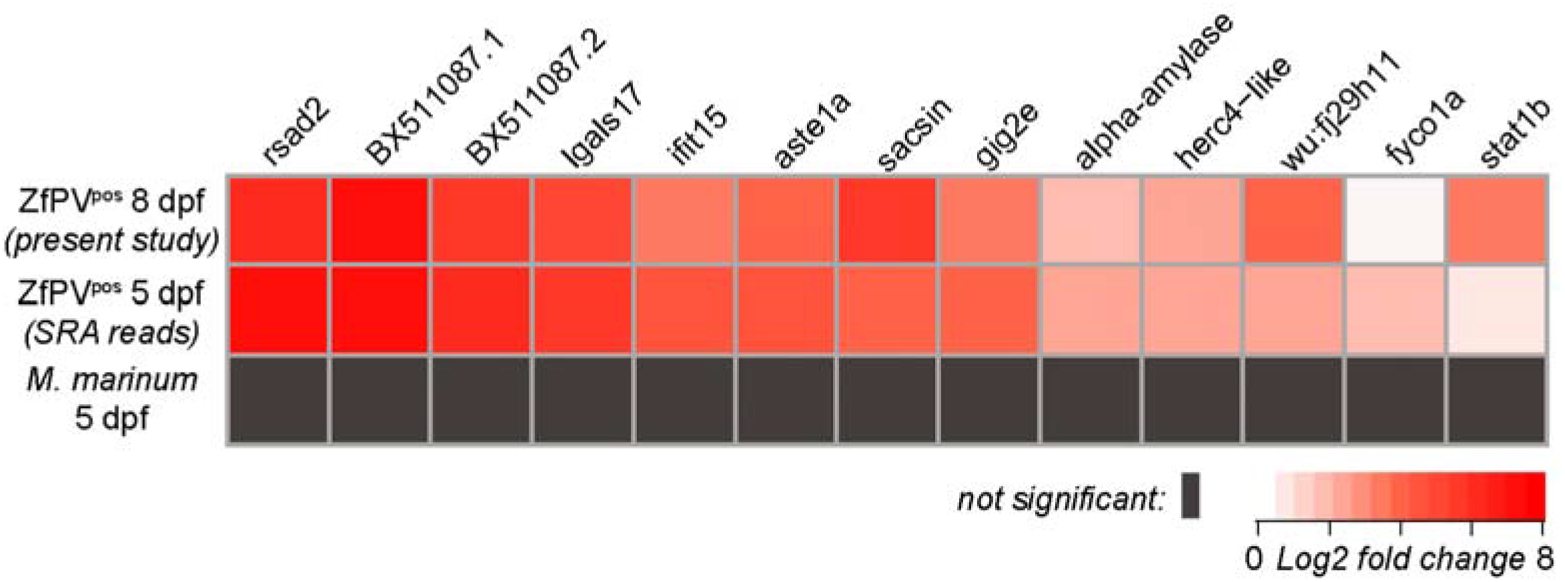
Genes induced by ZfPV in 5dpf and 8dpf zebrafish are not significantly induced by *Mycobacterium marinum* infection. Differential gene expression between WT animals infected with *M. marinum* and control animals (Yang et al., 2019) was quantified with reads from BioProject PRJNA398644. None of the genes induced by ZfPV infection in 5dpf and 8dpf were significantly induced by *M. marinum* infection (adjusted p-values of less than 0.1). Genes listed here include all genes that were significantly induced by ZfPV in 5dpf animals.

**Video 1.** Time-lapse recording in real time of an *isg15:GFP* animal at 8dpf that was treated with interferon as in Figure 1. The region of the animal captured in the video is the trunk vasculature. GFP-positive cells are apparent in circulation. Images were acquired with a Nikon Eclipse Ti Inverted Microscope with a VC Plan Apo 20X NA 0.75 objective (video file available upon request).

## References

Altan E, Kubiski SV, Boros A, Reuter G, Sadeghi M, Deng X, Creighton EK, Crim MJ, Delwart E. 2019. A Highly Divergent Picornavirus Infecting the Gut Epithelia of Zebrafish (Danio rerio) in Research Institutions Worldwide. Zebrafish 16:291–299. doi:10.1089/zeb.2018.1710

Bates JM, Mittge E, Kuhlman J, Baden KN, Cheesman SE, Guillemin K. 2006. Distinct signals from the microbiota promote different aspects of zebrafish gut differentiation. Dev Biol 297:374–386. doi:10.1016/j.ydbio.2006.05.006

Bermudez R, Losada AP, de Azevedo AM, Guerra-Varela J, Perez-Fernandez D, Sanchez L, Padros F, Nowak B, Quiroga MI. 2018. First description of a natural infection with spleen and kidney necrosis virus in zebrafish. J Fish Dis. doi:10.1111/jfd.12822

Binesh CP. 2013. Mortality due to viral nervous necrosis in zebrafish Danio rerio and goldfish Carassius auratus. Dis Aquat Organ 104:257–260. doi:10.3354/dao02605

Bouckaert R, Vaughan TG, Barido-Sottani J, Duchene S, Fourment M, Gavryushkina A, Heled J, Jones G, Kuhnert D, De Maio N, Matschiner M, Mendes FK, Muller NF, Ogilvie HA, du Plessis L, Popinga A, Rambaut A, Rasmussen D, Siveroni I, Suchard MA, Wu C-H, Xie D, Zhang C, Stadler T, Drummond AJ. 2019. BEAST 2.5: An advanced software platform for Bayesian evolutionary analysis. PLoS Comput Biol 15:e1006650. doi:10.1371/journal.pcbi.1006650

Cobian Guemes AG, Youle M, Cantu VA, Felts B, Nulton J, Rohwer F. 2016. Viruses as Winners in the Game of Life. Annu Rev Virol 3:197–214. doi:10.1146/annurev-virology-100114-054952

Crim MJ, Riley LK. 2012. Viral diseases in zebrafish: what is known and unknown. ILAR J 53:135–143. doi:10.1093/ilar.53.2.135

Darriba D, Taboada GL, Doallo R, Posada D. 2011. ProtTest 3: fast selection of best-fit models of protein evolution. Bioinforma Oxf Engl 27:1164–1165. doi:10.1093/bioinformatics/btr088

Di Tommaso P, Moretti S, Xenarios I, Orobitg M, Montanyola A, Chang J-M, Taly J-F, Notredame C. 2011. T-Coffee: a web server for the multiple sequence alignment of protein and RNA sequences using structural information and homology extension. Nucleic Acids Res 39:W13–17. doi:10.1093/nar/gkr245

Dirscherl H, Yoder JA. 2015. A nonclassical MHC class I U lineage locus in zebrafish with a null haplotypic variant. Immunogenetics 67:501–513. doi:10.1007/s00251-015-0862-1

Djordjevic D, Kusumi K, Ho JWK. 2016. XGSA: A statistical method for cross-species gene set analysis. Bioinforma Oxf Engl 32:i620–i628. doi:10.1093/bioinformatics/btw428

Dobin A, Davis CA, Schlesinger F, Drenkow J, Zaleski C, Jha S, Batut P, Chaisson M, Gingeras TR. 2013. STAR: ultrafast universal RNA-seq aligner. Bioinforma Oxf Engl 29:15–21. doi:10.1093/bioinformatics/bts635

Felix M-A, Ashe A, Piffaretti J, Wu G, Nuez I, Belicard T, Jiang Y, Zhao G, Franz CJ, Goldstein LD, Sanroman M, Miska EA, Wang D. 2011. Natural and experimental infection of Caenorhabditis nematodes by novel viruses related to nodaviruses. PLoS Biol 9:e1000586. doi:10.1371/journal.pbio.1000586

Finkbeiner SR, Allred AF, Tarr PI, Klein EJ, Kirkwood CD, Wang D. 2008. Metagenomic analysis of human diarrhea: viral detection and discovery. PLoS Pathog 4:e1000011. doi:10.1371/journal.ppat.1000011

Gentry RR, Froehlich HE, Grimm D, Kareiva P, Parke M, Rust M, Gaines SD, Halpern BS. 2017. Mapping the global potential for marine aquaculture. Nat Ecol Evol 1:1317–1324. doi:10.1038/s41559-017-0257-9

Geoghegan JL, Di Giallonardo F, Cousins K, Shi M, Williamson JE, Holmes EC. 2018. Hidden diversity and evolution of viruses in market fish. Virus Evol 4:vey031. doi:10.1093/ve/vey031

Geoghegan JL, Duchêne S, Holmes EC. 2017. Comparative analysis estimates the relative frequencies of co-divergence and cross-species transmission within viral families. PLoS Pathog 13:e1006215. doi:10.1371/journal.ppat.1006215

Greninger AL, Runckel C, Chiu CY, Haggerty T, Parsonnet J, Ganem D, DeRisi JL. 2009. The complete genome of klassevirus - a novel picornavirus in pediatric stool. Virol J 6:82. doi:10.1186/1743-422X-6-82

Grimholt U, Larsen S, Nordmo R, Midtlyng P, Kjoeglum S, Storset A, Saebø S, Stet RJM. 2003. MHC polymorphism and disease resistance in Atlantic salmon (Salmo salar); facing pathogens with single expressed major histocompatibility class I and class II loci. Immunogenetics 55:210–219. doi:10.1007/s00251-003-0567-8

Gruber AR, Lorenz R, Bernhart SH, Neubock R, Hofacker IL. 2008. The Vienna RNA websuite. Nucleic Acids Res 36:W70–74. doi:10.1093/nar/gkn188

Guindon S, Dufayard J-F, Lefort V, Anisimova M, Hordijk W, Gascuel O. 2010. New algorithms and methods to estimate maximum-likelihood phylogenies: assessing the performance of PhyML 3.0. Syst Biol 59:307–321. doi:10.1093/sysbio/syq010

Haas BJ, Papanicolaou A, Yassour M, Grabherr M, Blood PD, Bowden J, Couger MB, Eccles D, Li B, Lieber M, MacManes MD, Ott M, Orvis J, Pochet N, Strozzi F, Weeks N, Westerman R, William T, Dewey CN, Henschel R, LeDuc RD, Friedman N, Regev A. 2013. De novo transcript sequence reconstruction from RNA-seq using the Trinity platform for reference generation and analysis. Nat Protoc 8:1494–1512. doi:10.1038/nprot.2013.084

Hamming OJ, Lutfalla G, Levraud J-P, Hartmann R. 2011. Crystal structure of Zebrafish interferons I and II reveals conservation of type I interferon structure in vertebrates. J Virol 85:8181–8187. doi:10.1128/JVI.00521-11

Hunt HD, Jadhao S, Swayne DE. 2010. Major histocompatibility complex and background genes in chickens influence susceptibility to high pathogenicity avian influenza virus. Avian Dis 54:572–575. doi:10.1637/8888-042409-ResNote.1

Kwan KM, Fujimoto E, Grabher C, Mangum BD, Hardy ME, Campbell DS, Parant JM, Yost HJ, Kanki JP, Chien C-B. 2007. The Tol2kit: a multisite gateway-based construction kit for Tol2 transposon transgenesis constructs. Dev Dyn Off Publ Am Assoc Anat 236:3088–3099. doi:10.1002/dvdy.21343

Langmead B, Salzberg SL. 2012. Fast gapped-read alignment with Bowtie 2. Nat Methods 9:357–359. doi:10.1038/nmeth.1923

Levraud J-P, Palha N, Langevin C, Boudinot P. 2014. Through the looking glass: witnessing host-virus interplay in zebrafish. Trends Microbiol 22:490–497. doi:10.1016/j.tim.2014.04.014

Li H, Handsaker B, Wysoker A, Fennell T, Ruan J, Homer N, Marth G, Abecasis G, Durbin R. 2009. The Sequence Alignment/Map format and SAMtools. Bioinforma Oxf Engl 25:2078–2079. doi:10.1093/bioinformatics/btp352

Li L, Victoria JG, Wang C, Jones M, Fellers GM, Kunz TH, Delwart E. 2010. Bat guano virome: predominance of dietary viruses from insects and plants plus novel mammalian viruses. J Virol 84:6955–6965. doi:10.1128/JVI.00501-10

Lienenklaus S, Cornitescu M, Zietara N, Lyszkiewicz M, Gekara N, Jablonska J, Edenhofer F, Rajewsky K, Bruder D, Hafner M, Staeheli P, Weiss S. 2009. Novel reporter mouse reveals constitutive and inflammatory expression of IFN-beta in vivo. J Immunol Baltim Md 1950 183:3229–3236. doi:10.4049/jimmunol.0804277

Lim ES, Cao S, Holtz LR, Antonio M, Stine OC, Wang D. 2014. Discovery of rosavirus 2, a novel variant of a rodent-associated picornavirus, in children from The Gambia. Virology 454–455:25–33. doi:10.1016/j.virol.2014.01.018

Love MI, Huber W, Anders S. 2014. Moderated estimation of fold change and dispersion for RNA-seq data with DESeq2. Genome Biol 15:550. doi:10.1186/s13059-014-0550-8

Lysholm F, Wetterbom A, Lindau C, Darban H, Bjerkner A, Fahlander K, Lindberg AM, Persson B, Allander T, Andersson B. 2012. Characterization of the viral microbiome in patients with severe lower respiratory tract infections, using metagenomic sequencing. PloS One 7:e30875. doi:10.1371/journal.pone.0030875

Matzaraki V, Kumar V, Wijmenga C, Zhernakova A. 2017. The MHC locus and genetic susceptibility to autoimmune and infectious diseases. Genome Biol 18:76. doi:10.1186/s13059-017-1207-1

McConnell SC, Hernandez KM, Wcisel DJ, Kettleborough RN, Stemple DL, Yoder JA, Andrade J, de Jong JLO. 2016. Alternative haplotypes of antigen processing genes in zebrafish diverged early in vertebrate evolution. Proc Natl Acad Sci U S A 113:E5014–5023. doi:10.1073/pnas.1607602113

McConnell SC, Restaino AC, de Jong JLO. 2014. Multiple divergent haplotypes express completely distinct sets of class I MHC genes in zebrafish. Immunogenetics 66:199–213. doi:10.1007/s00251-013-0749-y

McCurley AT, Callard GV. 2008. Characterization of housekeeping genes in zebrafish: male-female differences and effects of tissue type, developmental stage and chemical treatment. BMC Mol Biol 9:102. doi:10.1186/1471-2199-9-102

Mistry J, Finn RD, Eddy SR, Bateman A, Punta M. 2013. Challenges in homology search: HMMER3 and convergent evolution of coiled-coil regions. Nucleic Acids Res 41:e121. doi:10.1093/nar/gkt263

Mizgirev IV, Revskoy S. 2010. A new zebrafish model for experimental leukemia therapy. Cancer Biol Ther 9:895–902. doi:10.4161/cbt.9.11.11667

Olival KJ, Hosseini PR, Zambrana-Torrelio C, Ross N, Bogich TL, Daszak P. 2017. Host and viral traits predict zoonotic spillover from mammals. Nature 546:646–650. doi:10.1038/nature22975

Palha N, Guivel-Benhassine F, Briolat V, Lutfalla G, Sourisseau M, Ellett F, Wang C-H, Lieschke GJ, Herbomel P, Schwartz O, Levraud J-P. 2013. Real-time wholebody visualization of Chikungunya Virus infection and host interferon response in zebrafish. PLoS Pathog 9:e1003619. doi:10.1371/journal.ppat.1003619

Parks GD, Duke GM, Palmenberg AC. 1986. Encephalomyocarditis virus 3C protease: efficient cell-free expression from clones which link viral 5’ noncoding sequences to the P3 region. J Virol 60:376–384.

Patowary A, Purkanti R, Singh M, Chauhan R, Singh AR, Swarnkar M, Singh N, Pandey V, Torroja C, Clark MD, Kocher J-P, Clark KJ, Stemple DL, Klee EW, Ekker SC, Scaria V, Sivasubbu S. 2013. A sequence-based variation map of zebrafish. Zebrafish 10:15–20. doi:10.1089/zeb.2012.0848

Patro R, Duggal G, Love MI, Irizarry RA, Kingsford C. 2017. Salmon provides fast and bias-aware quantification of transcript expression. Nat Methods 14:417–419. doi:10.1038/nmeth.4197

Phelan PE, Pressley ME, Witten PE, Mellon MT, Blake S, Kim CH. 2005. Characterization of snakehead rhabdovirus infection in zebrafish (Danio rerio). J Virol 79:1842–1852. doi:10.1128/JVI.79.3.1842-1852.2005

Racaniello VR. 2006. One hundred years of poliovirus pathogenesis. Virology 344:9–16. doi:10.1016/j.virol.2005.09.015

Rambaut A, Drummond AJ, Xie D, Baele G, Suchard MA. 2018. Posterior Summarization in Bayesian Phylogenetics Using Tracer 1.7. Syst Biol 67:901–904. doi:10.1093/sysbio/syy032

Robinson JT, Thorvaldsdottir H, Winckler W, Guttman M, Lander ES, Getz G, Mesirov JP. 2011. Integrative genomics viewer. Nat Biotechnol 29:24–26. doi:10.1038/nbt.1754

Roediger B, Lee Q, Tikoo S, Cobbin JCA, Henderson JM, Jormakka M, O’Rourke MB, Padula MP, Pinello N, Henry M, Wynne M, Santagostino SF, Brayton CF, Rasmussen L, Lisowski L, Tay SS, Harris DC, Bertram JF, Dowling JP, Bertolino P, Lai JH, Wu W, Bachovchin WW, Wong JJ-L, Gorrell MD, Shaban B, Holmes EC, Jolly CJ, Monette S, Weninger W. 2018. An Atypical Parvovirus Drives Chronic Tubulointerstitial Nephropathy and Kidney Fibrosis. Cell 175:530–543.e24. doi:10.1016/j.cell.2018.08.013

Roeselers G, Mittge EK, Stephens WZ, Parichy DM, Cavanaugh CM, Guillemin K, Rawls JF. 2011. Evidence for a core gut microbiota in the zebrafish. ISME J 5:1595–1608. doi:10.1038/ismej.2011.38

Ryan MD, King AM, Thomas GP. 1991. Cleavage of foot-and-mouth disease virus polyprotein is mediated by residues located within a 19 amino acid sequence. J Gen Virol 72 (Pt 11):2727–2732. doi:10.1099/0022-1317-72-11-2727

Sanders GE, Batts WN, Winton JR. 2003. Susceptibility of zebrafish (Danio rerio) to a model pathogen, spring viremia of carp virus. Comp Med 53:514–521.

Schmittgen TD, Livak KJ. 2008. Analyzing real-time PCR data by the comparative C(T) method. Nat Protoc 3:1101–1108.

Secombes CJ, Zou J. 2017. Evolution of Interferons and Interferon Receptors. Front Immunol 8:209. doi:10.3389/fimmu.2017.00209

Seppola M, Stenvik J, Steiro K, Solstad T, Robertsen B, Jensen I. 2007. Sequence and expression analysis of an interferon stimulated gene (ISG15) from Atlantic cod (Gadus morhua L.). Dev Comp Immunol 31:156–171. doi:10.1016/j.dci.2006.05.009

Sharma D, Sehgal P, Mathew S, Vellarikkal SK, Singh AR, Kapoor S, Jayarajan R, Scaria V, Sivasubbu S. 2019. A genome-wide map of circular RNAs in adult zebrafish. Sci Rep 9:3432. doi:10.1038/s41598-019-39977-7

Shi M, Lin X-D, Chen X, Tian J-H, Chen L-J, Li K, Wang W, Eden J-S, Shen J-J, Liu L, Holmes EC, Zhang Y-Z. 2018. The evolutionary history of vertebrate RNA viruses. Nature 556:197–202. doi:10.1038/s41586-018-0012-7

Shi M, Lin X-D, Tian J-H, Chen L-J, Chen X, Li C-X, Qin X-C, Li J, Cao J-P, Eden J-S, Buchmann J, Wang W, Xu J, Holmes EC, Zhang Y-Z. 2016. Redefining the invertebrate RNA virosphere. Nature 540:539–543. doi:10.1038/nature20167

Suttle CA. 2005. Viruses in the sea. Nature 437:356–361. doi:10.1038/nature04160

Swatek KN, Aumayr M, Pruneda JN, Visser LJ, Berryman S, Kueck AF, Geurink PP,Ovaa H, van Kuppeveld FJM, Tuthill TJ, Skern T, Komander D. 2018. Irreversible inactivation of ISG15 by a viral leader protease enables alternative infection detection strategies. Proc Natl Acad Sci U S A 115:2371–2376. doi:10.1073/pnas.1710617115

Tcherepanov V, Ehlers A, Upton C. 2006. Genome Annotation Transfer Utility (GATU): rapid annotation of viral genomes using a closely related reference genome. BMC Genomics 7:150. doi:10.1186/1471-2164-7-150

Tobin DM, May RC, Wheeler RT. 2012. Zebrafish: a see-through host and a fluorescent toolbox to probe host-pathogen interaction. PLoS Pathog 8:e1002349. doi:10.1371/journal.ppat.1002349

Van Dycke J, Ny A, Conceição-Neto N, Maes J, Hosmillo M, Cuvry A, Goodfellow I, Nogueira TC, Verbeken E, Matthijnssens J, de Witte P, Neyts J, Rocha-Pereira J. 2019. A robust human norovirus replication model in zebrafish larvae. PLoS Pathog 15:e1008009. doi:10.1371/journal.ppat.1008009

Varela M, Figueras A, Novoa B. 2017. Modelling viral infections using zebrafish: Innate immune response and antiviral research. Antiviral Res 139:59–68. doi:10.1016/j.antiviral.2016.12.013

Wallace KN, Akhter S, Smith EM, Lorent K, Pack M. 2005. Intestinal growth and differentiation in zebrafish. Mech Dev 122:157–173. doi:10.1016/j.mod.2004.10.009

Warren DL, Geneva AJ, Lanfear R. 2017. RWTY (R We There Yet): An R Package for Examining Convergence of Bayesian Phylogenetic Analyses. Mol Biol Evol 34:1016–1020. doi:10.1093/molbev/msw279

Webster CL, Waldron FM, Robertson S, Crowson D, Ferrari G, Quintana JF, Brouqui J-M, Bayne EH, Longdon B, Buck AH, Lazzaro BP, Akorli J, Haddrill PR, Obbard DJ. 2015. The Discovery, Distribution, and Evolution of Viruses Associated with Drosophila melanogaster. PLoS Biol 13:e1002210. doi:10.1371/journal.pbio.1002210

Westerfield M. 2007. The Zebrafish Book: A guide for the laboratory use of zebrafish (Danio rerio), 5th ed. Eugene: University of Oregon Press.

White RJ, Collins JE, Sealy IM, Wali N, Dooley CM, Digby Z, Stemple DL, Murphy DN, Billis K, Hourlier T, Fullgrabe A, Davis MP, Enright AJ, Busch-Nentwich EM. 2017. A high-resolution mRNA expression time course of embryonic development in zebrafish. eLife 6. doi:10.7554/eLife.30860

Yang S, Hu W, Shimada Y, Münch M, Marín-Juez R, Meijer AH, Spaink HP. 2019. RNA-seq analysis of a zebrafish tlr2 mutant shows a broad function of this Tolllike receptor in transcriptional and metabolic control and defense to Mycobacterium marinum infection. bioRxiv 742601. doi:10.1101/742601

Zell R. 2018. Picornaviridae-the ever-growing virus family. Arch Virol 163:299–317. doi:10.1007/s00705-017-3614-8

Zell R, Delwart E, Gorbalenya AE, Hovi T, King AMQ, Knowles NJ, Lindberg AM, Pallansch MA, Palmenberg AC, Reuter G, Simmonds P, Skern T, Stanway G, Yamashita T, Ictv Report Consortium null. 2017. ICTV Virus Taxonomy Profile: Picornaviridae. J Gen Virol 98:2421–2422. doi:10.1099/jgv.0.000911

Zhao J, Wu J, Xu T, Yang Q, He J, Song X. 2018. IRESfinder: Identifying RNA internal ribosome entry site in eukaryotic cell using framed k-mer features. J Genet Genomics Yi Chuan Xue Bao 45:403–406. doi:10.1016/j.jgg.2018.07.006

